# Design and Computational Analysis of a Chimeric Avian Influenza Antigen: A Yeast-displayed, Universal and Cross-protective Vaccine Candidate

**DOI:** 10.1101/2021.09.05.459052

**Authors:** Elyas Mohammadi, Zana Pirkhezranian, Samira Dashty, Naghmeh Saedi, Mohammad Hadi Sekhavati

## Abstract

**Background:** A cross-protective avian influenza vaccine candidate can be designed by using a preserved antigen against mutation in various subtypes of influenza. M2e peptide sequence has remained remarkably unchanged in influenza type A isolated since 1918.

**Methods:** A consensus sequence of M2e peptide was obtained from 31 sequences of H5N8, H5N1, H9N2 and H7N9 subtypes of avian influenza virus isolated from 7 avian species in 5 Asian countries. A partial sequence of flagellin was considered as an adjuvant. Subsequently, two chimeric antigens were designed to be virtually cloned and expressed using PYD1 vector and *EBY100* yeast strain. The stability and conformational features of these two antigens were assessed through molecular dynamic (MD) simulations. The detectability of vaccine candidates by a specific monoclonal antibody (MAb148) were estimated through docking studies.

**Results:** In spite of significant compactness and stability of the first candidate in comparison with the second design, it was less detectable by MAb148. Contrary to the first chimeric antigen, Van der Waals, electrostatic and binding energies of the interaction of the second antigen with MAb148 were significantly closer to the positive control. It is shown that epitopes of the second chimeric antigen could be correctly located in the specific pocket of CDR region of MAb148.

**Conclusion:** The second chimeric antigen could be considered as a yeast-displayed avian influenza vaccine candidate due to the capability of provoking humoral immunity and innate immune system by M2e and flagellin respectively.

## Introduction

Influenza virus consists of a single-stranded genomic fragment originated from the *orthomyxoviridae* family. At the moment, this family is categorized to 5 types of influenza: A, B, C, *Thogotovirus* and *Isavirus*. Among these categories, only type A can be pathogenic in birds and it is classified to various subtypes with different pathogenicity levels according to genetic variation in surface glycoproteins of hemagglutinin (HA) (16 HA) and neuraminidase (NA) (9 NA)(1). Due to significant genetic diversity even between the same viruses of a pandemic subtype, these influenza viruses encounter with a much weaker immune response in the first infection. Therefore, prevalence of pandemic influenza can cause disease, death and economic losses in avian industry (2). Antiviral drugs against influenza are available in the market (3, 4) but resistance of the virus to treatments can grow significantly. Between 2008 and 2009, nearly 100% of seasonal influenza H1N1 and H3N2 subtypes in the United States were unsusceptible to Oseltamivir and Zanamivir (two commercially available drugs against Influenza), respectively.

Efforts to produce a recombinant vaccine using HA and NA as two major antigens of influenza virus have raised many problems because of genetic variability (3, 5, 6). Vaccination with a conserve antigen against mutations can create immunity against several subtypes of influenza (7, 8). In this regard, protein Matrix 2 (M2) can be considered as the most substantial antigen. The outer membrane section of this antigen named M2e is exposed and could be recognized by immune system. This segment consists of 24 amino acids and it forms an ion channel which plays a vital role in virus replication (9). Nine amino acids of this segment are remarkably conserved among all subtypes of influenza (9-11). This section is completely identical in 1364 sequences extracted from NCBI database (12). “SLLTE” sequence as the main core which brings antigenicity of M2e has 97%, 98%, and 98% identity in human, swine and avian M2e sequences, respectively (12).

In spite of lower antigenicity of M2e peptide in comparison with HA and NA, it is considered as a target for vaccine design due to being remained unchanged in all of influenza type A isolated since 1918 (9, 13). Moreover, expression of this peptide is two times more than HA on the surface of infected cells (14, 15) which may overcome its lower antigenicity (16). For vaccine design, using M2e antigen individually may provoke immune system weakly, but studies have shown that multiple usage of this peptide sequence increase the immunogenic effects (17-20). *In vivo* studies have revealed that specific antibodies against M2e peptide can diminish the lesions or fatality consequences of several subtypes of influenza(14, 21-23). in order to increase the antigenicity of M2e I previous studies, it was fused to various adjuvants or carriers which lead to enhancing protection against lethal challenges (24). Many investigations have also shown that M2e-based vaccines can generate immunity against different influenza subtypes (10, 17, 19, 25-27).

Additionally, it has been suggested that M2e-based vaccines can be considered as a vaccine supplement to increase cross-protection ability (24, 28). Live attenuated influenza vaccine in combination with M2e Virus-like particles (VLPs) could protect mice against H3N2, H1N1 and H5N1 lethal challenges (28).

Flagellin, a principal component of bacterial flagella, is essential for bacterial movements. Flagellin is considered as one of the most important targets of immune system and acts as a ligand to active Toll-Like Receptor 5 (TLR5) in host cells (29-33). Stimulation of TLR5 leads to the activation of innate immune system (34). A chimeric protein consists of flagellin and a specific antigen has been used in vaccine development against variety of infections such as Neil, Malaria, plague and Tuberculosis (35-38). Several studies have pointed to the applications of flagellin as an effective adjuvant (29, 34). Other advantages of flagellin that make it a good candidate for vaccine production are: a) it is effective in low doses (39), b) it does not cause an IgE response (40), c) the presence of previous immunity to flagellin does not interfere with its function as an adjuvant (40, 41), d) no toxicity was detected in rabbits by nasal and intramuscular consumption and e) it can be easily produced in large quantities (34).

Recently, yeast-displayed vaccines have been studied for influenza and the results have shown that they can be advantageous due to the ability of large-scale production (42-47) and no necessity to add adjuvants like aluminum (44). Compared to the expression of virus proteins in a soluble form in the media, the expression of these proteins at the cell surface make them significantly more immunogenic and detectable by the host immune system (42, 48). In light of the above facts, in this study we designed two yeast-displayed chimeric avian influenza antigens consist of M2e peptide and flagellin. Finally, using computational-based approaches the fidelity and immunogenicity of these chimeric antigens were assessed.

## Material and methods

### Determining the consensus sequence of M2e antigen from four subtypes of avian influenza virus

Consensus sequence was provided based on a protocol conducted by Huleatt et al. in 2008 and Mozdzanowska et al. in 2003. To this end, 31 sequence of M2e protein isolated from 7 avian species which originated from 5 Asian countries (China, Japan, India, South Korea and Vietnam) retrieved from NCBI database (https://www.ncbi.nlm.nih.gov). Alignment process was done subsequently by CLC workbench 5 software to achieve consensus sequence of this antigen from H5N8, H5N1, H9N2 and H7N9 subtype of avian influenza (Table 1).

**Table 1.**
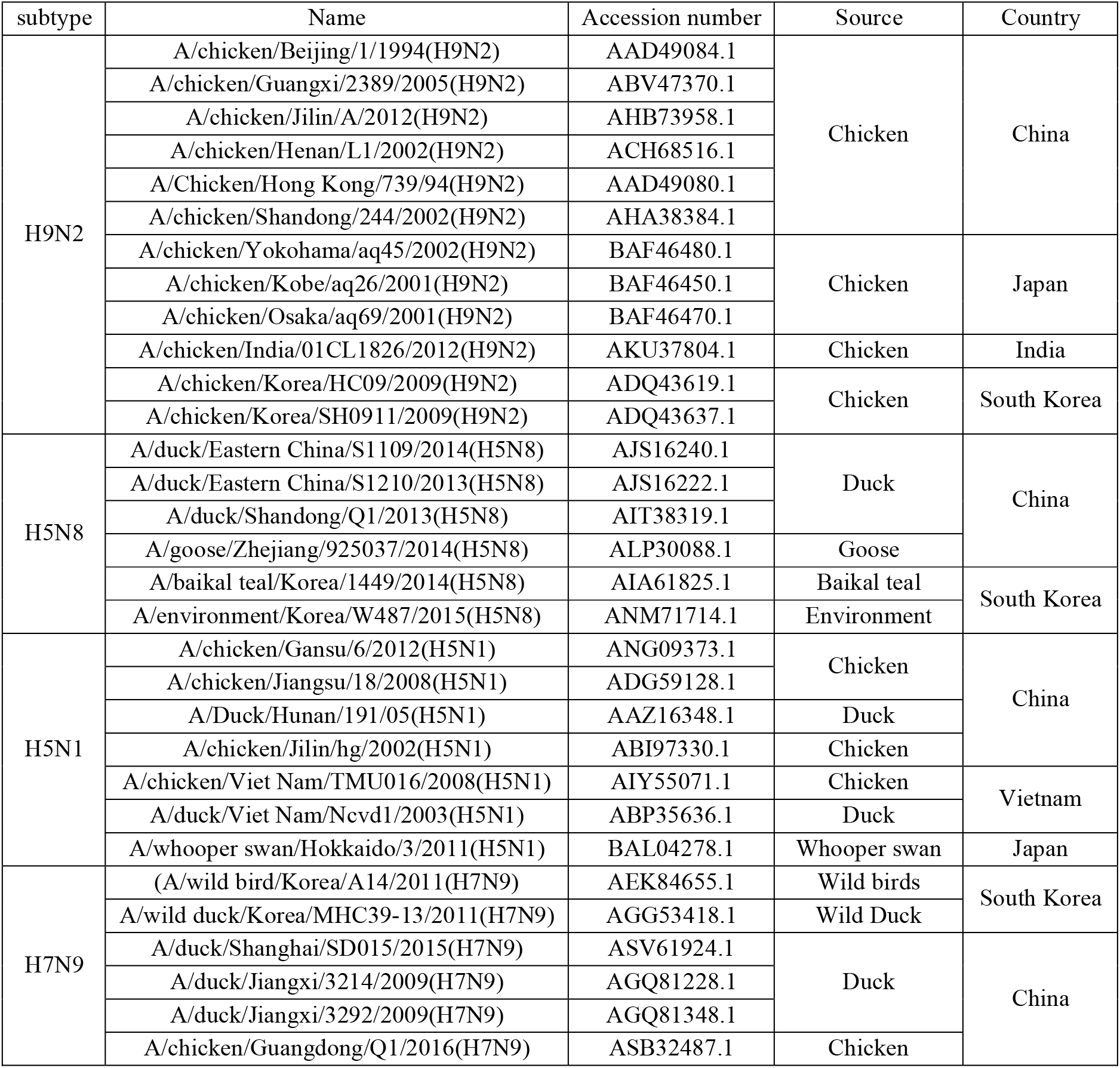
31 M2e sequences of 4 subtypes of influenza from 7 avian species which were originated from 5 Asian countries.

### Design of the chimeric antigens

In order to design the chimeric antigens, previous studies (16, 19, 34, 49-51) and information related to PYD1 shuttle vector (Addgene plasmid # 73447; http://n2t.net/addgene:73447; RRID: Addgene_73447) (52) and *EBY100* yeast strain (the *EGY100* yeast strain is genetically modified and contains the plasmid, pIU211 stably integrated into the genome for yeast-displaying proteins(53)) were used. In this regard, after obtaining the consensus sequence of M2e protein from H9N2, H5N1, H5N8 and H7N9 subtypes of avian influenza virus, a partial sequence of flagellin (retrieved from 5GY2, a TLR5-flagellin complex) was selected as an adjuvant to be fused to the consensus sequence of M2e (54). The Aga2 protein which is expressed by PYD1 vector, is specifically bound to Aga1 protein on the surface of *EBY100* yeast strain. Two flexible linkers were considered between Aga2 and next protein (ASGGGGSGGGGSGGGGS) and also between flagellin and 4 tandem copies of M2e antigen. According to this information, two different chimeric proteins were designed for displaying on the surface of *EBY100* yeast strain (Figure 1,A).

**Figure 1.**
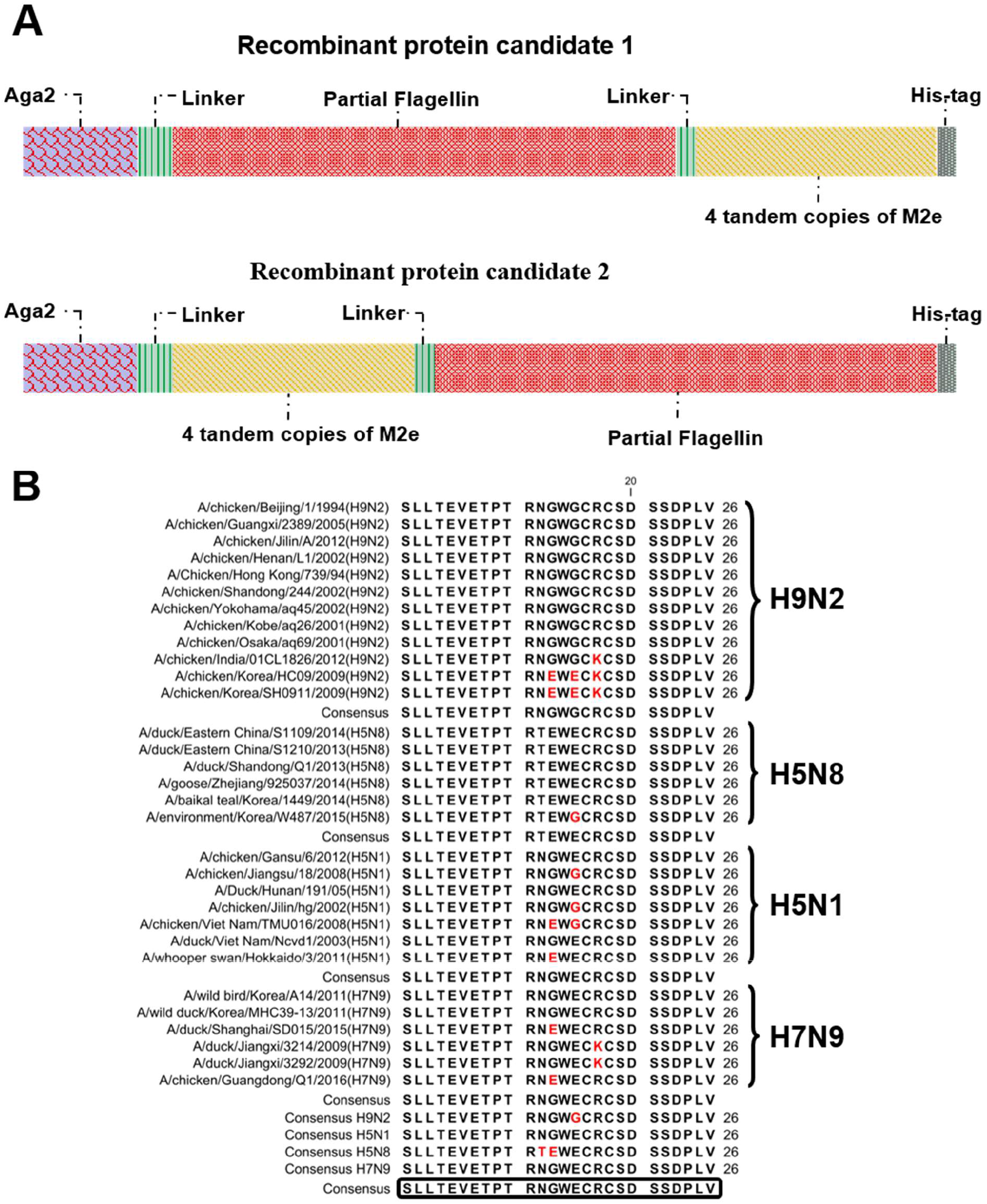
**A)** The schematic representation of two candidates of chimeric antigens from H5N8, H5N1, H9N2 and H7N9 subtypes of avian influenza virus. **Aga2:** a protein which could be expressed by PYD1 vector for binding to Aga1 protein on the surface of *EBY100* yeast strain, **flexible linker:** ASGGGGSGGGGSGGGGS, **4 tandem copies of M2e:** (SLLTEVETPTRNGWECRCSDSSDPLV)_4_, **partial sequence of flagellin:** it is considered as an adjutant, **His-tag:** for purification purposes. **B)** The consensus sequence of M2e protein from 4 subtypes (H9N9, H5N8, H5N1 and H9N2) of avian influenza virus and 7 avian species that originated from Asian countries (China, Japan, India, South Korea and Vietnam) were proposed.

### Protein Modeling

In order to investigate the function of designed chimeric antigens, first, the 3-dimensional (3D) structure of these peptides should be predicted. In this regard, the structure of designed proteins were predicted through I-TASSER (55) server. In accordance with previous studies, we substitute the cysteine with serine in 4 replications of M2e protein which will lead to the prevention of undesired disulfide bonds (SLLTEVETPTRNGWE<u>S</u>R<u>S</u>SDSSDPLV). Serine and cysteine displacement would not change the protein’s antigenic potency (16, 19). In order to model the designed proteins by I-TASSER, chain D of the 5GY2 crystallography structure (flagellin protein attached to the TLR5 receptor) was used. In addition, according to the crystallography structure of 5DLM (a section of the M2e protein which binds to the monoclonal antibody), the loop structure was applied to model the “SLLTEVETP” epitope.

### Molecular dynamic simulations

Molecular dynamic simulations were used to predict the structure of designed proteins in avian physiological condition (avian normal body temperature and pressure). All simulations were carried out using GROMACS 5 program (42, 56-58). Two designed proteins and docked complexes in further steps (protein-antibody) were processed under GROMOS 54a7 (59) force field library. Besides, SPC water model (60, 61) was used for the solvation in a periodic cubic box that was large enough to contain the system and 1 nm of solvent on all sides. For the neutralization of solvated complex, combination of Na^+^ and Cl^-^ were used. Neutral systems were then subjected to the steepest descent energy minimization. After energy minimization each system was equilibrated for 200ps under NVT and NPT conditions. Temperature was set to 313K. Final simulation was carried at NPT condition without any restraints. Pressure and temperature of the system were controlled by the Parrinello-Rahman (62) and V-rescale (63) algorithms respectively. The LINCS algorithm (64) was used to constrain all the bond lengths. A Verlet cutoff method was used for nonbonded interactions. Nonbonded interactions within 1 nm were updated every 20 steps. Trajectories were analyzed with the help of VMD (65) and Xmgrace. Using g_mmpbsa tool (66), van der waals, electrostatic and binding energies for the interaction of designed proteins and complement-determining regions (CDR) of specific monoclonal antibody (in docking studies) were calculated. The same calculation was done for a positive control (5DLM, crystallography of naturalizing antibody and desired epitope) through g_mmpbsa tool for further assessments.

### Docking studies

Prior to docking studies, the 3D structure of designed recombinant antigens which were prepared through protein modeling and MD simulations, were refined by ReFOLD server (67) to correct residues in disallowed regions. The accuracy of the predicted models prior and after refinement were assessed by Ramachandran plot analysis. Afterward, refined models were used to investigate the detectability of their epitopes by CDR region of a specific monoclonal antibody (MAB148, retrieved from 5DLM complex). Docking studies performed by antibody mode of Cluspro server (68). Subsequently, the position of epitopes in the specific pocket of antibody was assessed and visualized through Pymol 1.8 software (69).

## Results

### Determining the consensus sequence of M2e antigen

According to the conservation of M2e protein in different subtypes of avian influenza, few differences within and between four subtypes were observed. Accordingly, “SLLTEVETPTRNGWECRCSDSSDPLV” sequence was proposed as a consensus sequence of M2e antigen for H9N2, H5N1, H5N8 and H7N9 subtypes of avian influenza (Figure 1,B).

### Protein modeling and dynamic simulations

After determining the consensus sequence of M2e protein from 4 subtypes of avian influenza virus, two designed proteins (Figure 1,A) were modeled by I-Tasser server. The structural stability of recombinant proteins were investigated through GROMACS during 100 nanosecond (ns). Accordingly, with regard to RMSD plot (Figure 2, B), both recombinant proteins become stable after 20ns. In spite of several fluctuations in RMSD plot of both designs, changes were reported less than 0.2 Angstrom.

**Figure 2.**
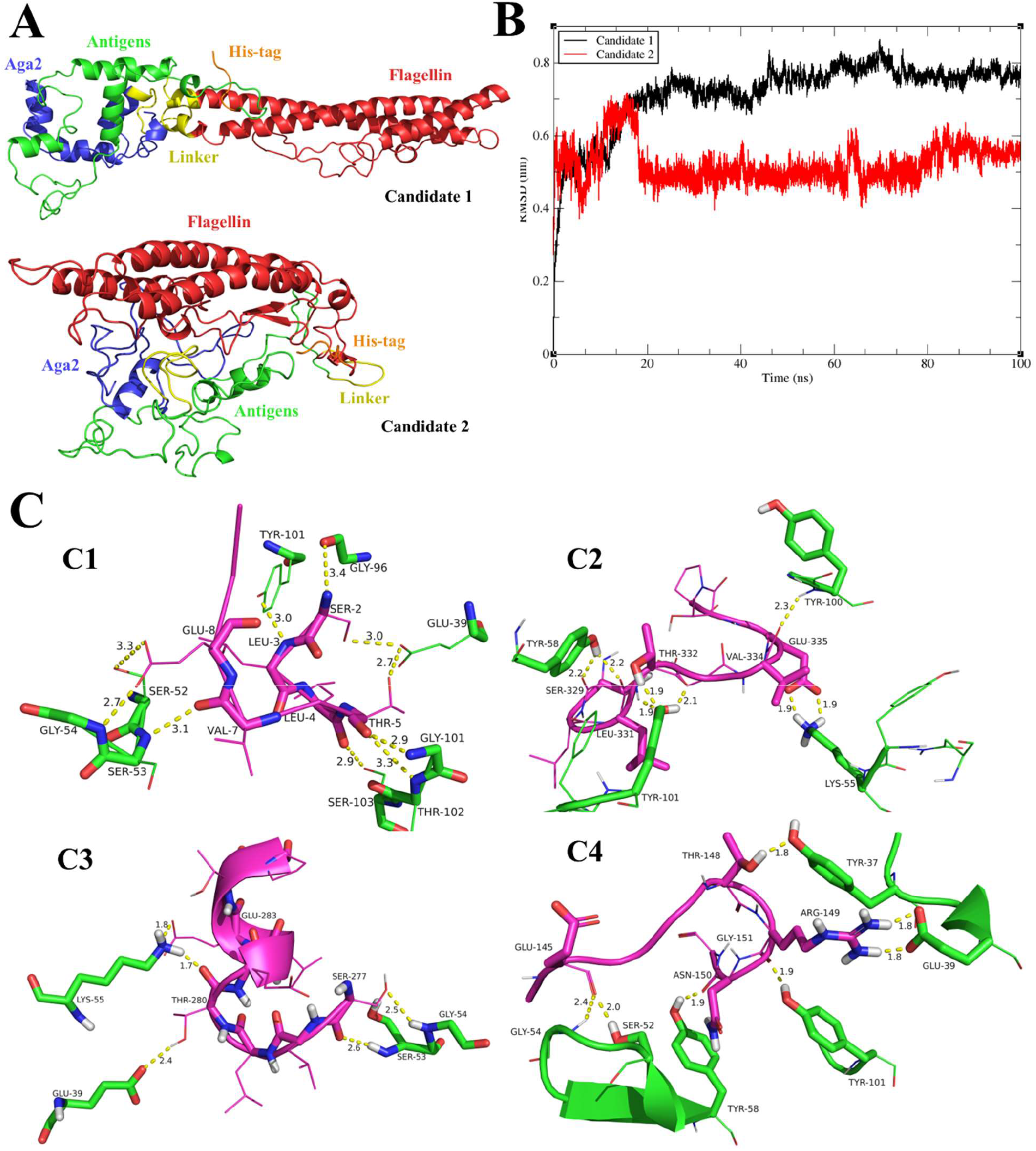
**A)** 3D structure of 2 proposed designs as yeast-displayed candidates after 100 nanoseconds MD simulations. **B)** The RMSD plot of both proteins based on alpha carbon space variation during the MD simulations. **C)** Cartoon representation of involved residues in docking of monoclonal antibody (green) and M2e protein epitopes (pink). **c1)** Crystallography complex of monoclonal antibody and epitope (positive control) **c2)** 1_329-337_ epitope and specific monoclonal antibody **c3)** 1_277-285_ epitope and specific monoclonal antibody **c4)** 2_143-151_ epitope and specific monoclonal antibody.

In terms of the first candidate, structural compactness of recombinant protein is less than the second candidate (Figure 2, A). The flagellin region is completely distanced from other parts and the antigenic section lacks any space barrier for binding to specific antibodies and also Aga2 protein is free for binding to the yeast surface.

On the other hand, the structural compactness of the second candidate almost caused inaccessibility of different sections, whereas this feature can play a vital role in the stability of protein. This fact may diminish the probability of antigenic recognition by specific antibodies which is investigated by docking analysis (Table 3).

### Docking and binding energy

Ramachandran plot analysis, prior and after structural refinement for the chimeric antigen models revealed significant changes for residues in disallowed regions (Table 2). The results of docking studies for two refined chimeric antigens indicated that two out of four desired epitopes (“SLLTEVETP”) in the first vaccine candidate is completely recognizable by CDR region of specific monoclonal antibody, whereas only a section of one desired epitope is detectable in the second one. According to g_mmpbsa tool results, despite the stronger Van der Waal energy between _143_EVETPTRNG_151_ epitope in the second candidate and its specific antibody in comparison with the two _329_SLLTEVETP_337_ and _277_SLLTEVETP_285_ epitopes in candidate 1, the electrostatic and binding energy of the identifiable epitopes of candidate 1 are significantly higher. Overall, the reported energies for epitopes of candidate 1 chimeric protein have the closest values to the positive control (binding, electrostatic, and Van der Waal energies between the “SLLTEVETP” epitope and CDR region of specific monoclonal antibody in crystallography structure, 5DLM) (Table 3).

**Table 2.**
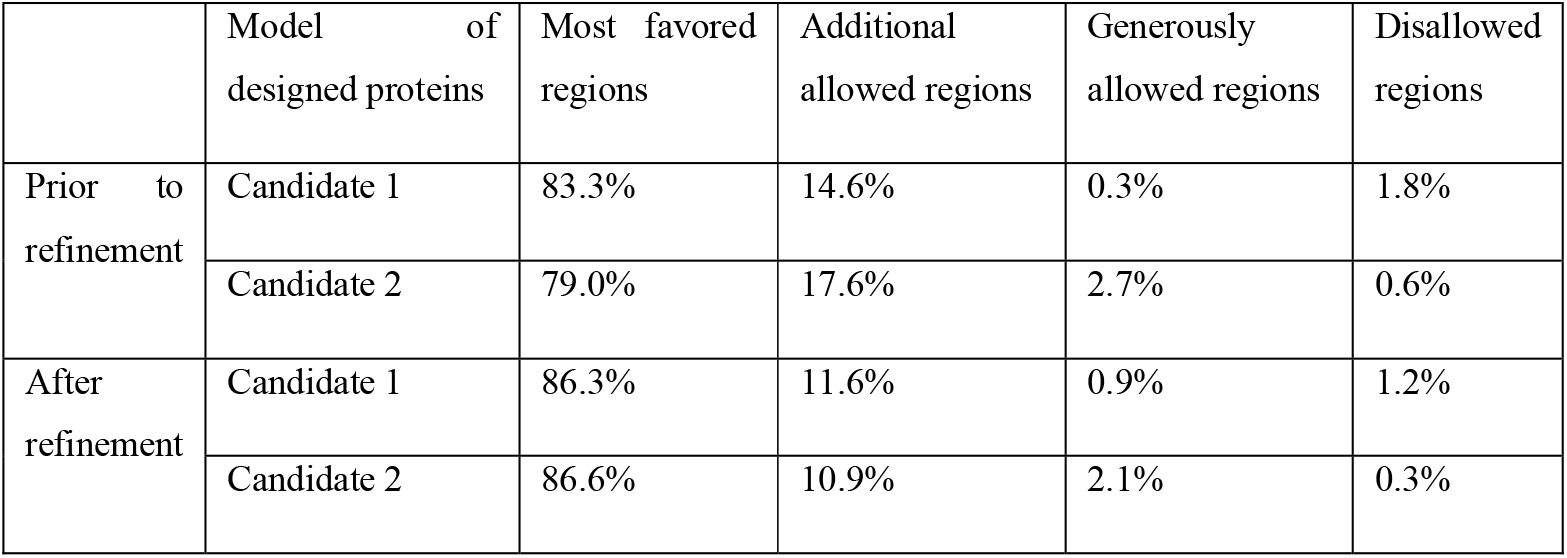
retrieved data from Ramachandran plot of designed proteins, prior to refinement and afterward.

|  | Model of designed proteins | Most favored regions | Additional allowed regions | Generously allowed regions | Disallowed regions |
| --- | --- | --- | --- | --- | --- |
| Prior to refinement | Candidate 1 | 83.3% | 14.6% | 0.3% | 1.8% |
|  | Candidate 2 | 79.0% | 17.6% | 2.7% | 0.6% |
| After refinement | Candidate 1 | 86.3% | 11.6% | 0.9% | 1.2% |
|  | Candidate 2 | 86.6% | 10.9% | 2.1% | 0.3% |

With regard to the residue involvement in antibody-epitope docking results, the _329_SLLTEVETP_337_ and _277_SLLTEVETP_285_ epitopes, which belong to the candidate 1 chimeric protein, were more similar to positive control (Table 3) (Figure 3,C).

**Table 3.**
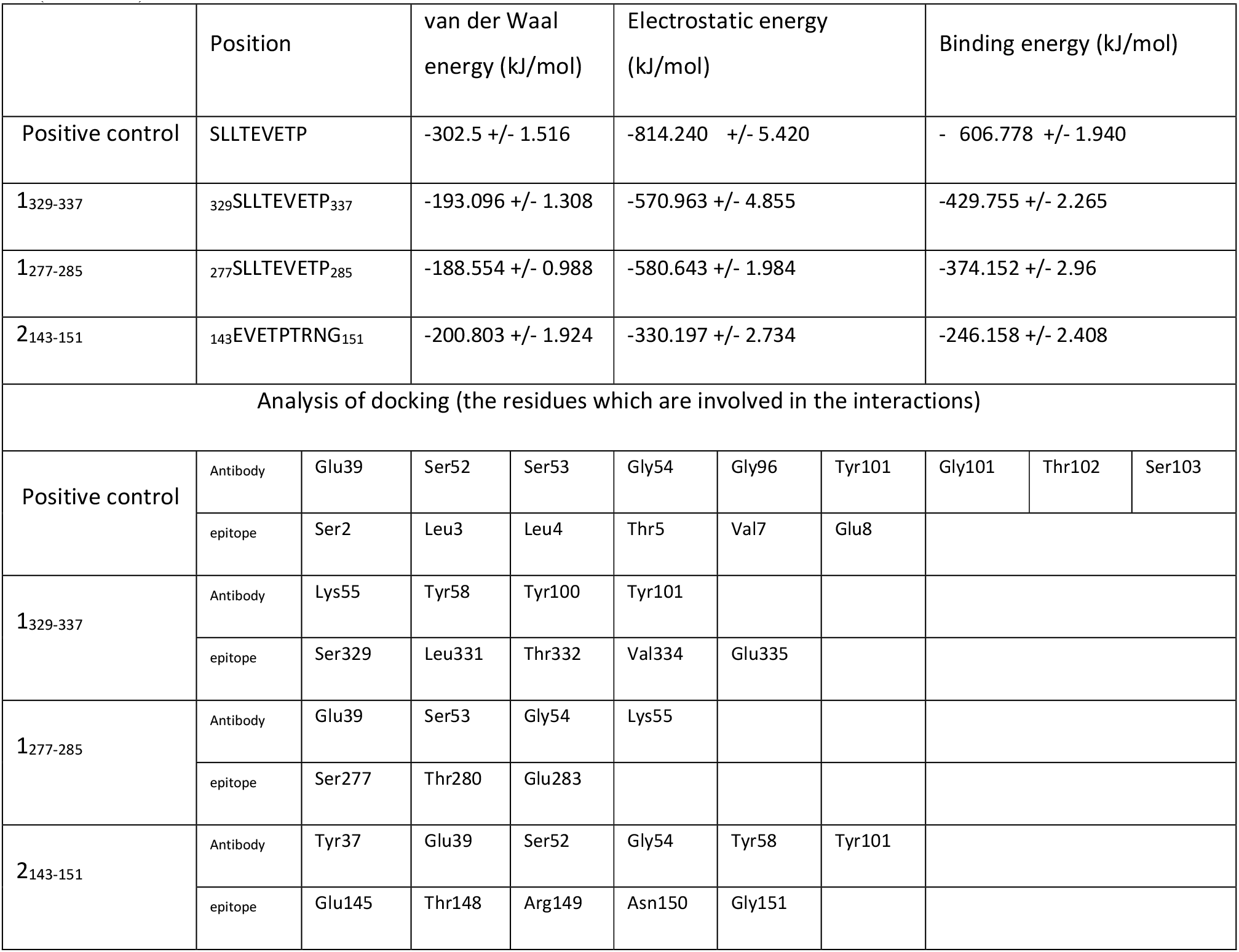
Van der Waal, electrostatics and binding energies and also involved residues in the interaction of “SLLTEVETP” epitope in positive control, candidate 1, and candidate 2 chimeric protein with the CDR region of the specific monoclonal antibody. **Positive control:** The “SLLTEVETP” epitope and CDR region of specific monoclonal antibody in crystallography structure (ID: 5DLM). **1**_**329-337**_ **and 1**_**277-285**_: First and second detectable epitope from candidate 1 chimeric protein by CDR region of specific monoclonal antibody (MAB148). **2**_**143-151**_: the only detectable epitope from candidate 2 chimeric protein by CDR region of specific monoclonal antibody (MAB148).

|  | Position | van der Waal<br>energy (kJ/mol) | Electrostatic energy<br>(kJ/mol) | Binding energy (kJ/mol) |  |  |  |  |  |  |
| --- | --- | --- | --- | --- | --- | --- | --- | --- | --- | --- |
| Positive control | SLLTEVETP | -302.5 +/- 1.516 | -814.240 +/- 5.420 | - 606.778 +/- 1.940 |  |  |  |  |  |  |
| 1<sub>329-337</sub> | 329SLLTEVETP<sub>337</sub> | -193.096 +/- 1.308 | -570.963 +/- 4.855 | -429.755 +/- 2.265 |  |  |  |  |  |  |
| 1<sub>277-285</sub> | 277SLLTEVETP<sub>285</sub> | -188.554 +/- 0.988 | -580.643 +/- 1.984 | -374.152 +/- 2.96 |  |  |  |  |  |  |
| 2<sub>143-151</sub> | 143EVETPTRNG<sub>151</sub> | -200.803 +/- 1.924 | -330.197 +/- 2.734 | -246.158 +/- 2.408 |  |  |  |  |  |  |
| Analysis of docking (the residues which are involved in the interactions) |  |  |  |  |  |  |  |  |  |  |
| Positive control | Antibody | Glu39 | Ser52 | Ser53 | Gly54 | Gly96 | Tyr101 | Gly101 | Thr102 | Ser103 |
|  | epitope | Ser2 | Leu3 | Leu4 | Thr5 | Val7 | Glu8 |  |  |  |
| 1<sub>329-337</sub> | Antibody | Lys55 | Tyr58 | Tyr100 | Tyr101 |  |  |  |  |  |
|  | epitope | Ser329 | Leu331 | Thr332 | Val334 | Glu335 |  |  |  |  |
| 1<sub>277-285</sub> | Antibody | Glu39 | Ser53 | Gly54 | Lys55 |  |  |  |  |  |
|  | epitope | Ser277 | Thr280 | Glu283 |  |  |  |  |  |  |
| 2<sub>143-151</sub> | Antibody | Tyr37 | Glu39 | Ser52 | Gly54 | Tyr58 | Tyr101 |  |  |  |
|  | epitope | Glu145 | Thr148 | Arg149 | Asn150 | Gly151 |  |  |  |  |

**Figure 3.**
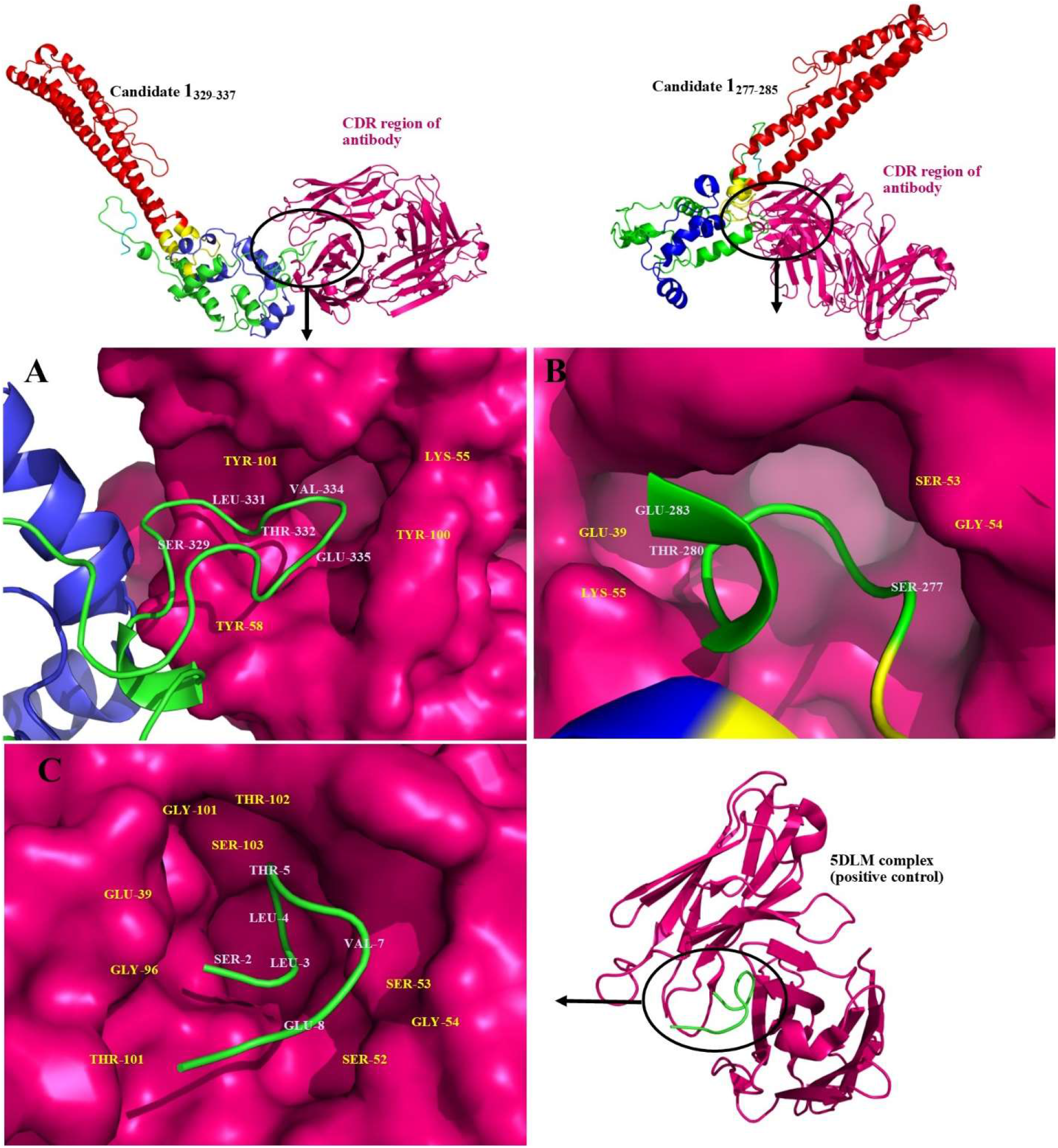
The position of 1_329-337_ and 1_277-285_ epitopes (first candidate) and “SLLTEVETP” epitope of crystallography (ID: 5DLM) in the specific pocket of CDR region of monoclonal antibody (the residues of antibody and epitope which involved in interactions are shown in yellow and white respectively). **A)** epitope 1_329-337_. **B)** epitope 1_277-285_. **C)** 5DLM complex (positive control).

In terms of docking results, the location of two desire epitopes of the first candidate in CDR region of a specific monoclonal antibody depict the accurate positioning of these two epitopes in the relevant pocket of monoclonal antibody (Figure 3). This fact can boost humoral immunity accurately and specifically.

### Virtual cloning

According to the results of docking studies and molecular dynamic simulations, and also by considering the structural properties of two candidate proteins, it can be concluded that protein candidate 1 chimeric protein is a better choice for expression on the yeast surface (Figure 4). Therefore, the coding sequence of this protein can be selected for expression on the yeast surface of *Saccharomyces cerevisiae (EBY100)*. By flanking the coding sequence of the first candidate between *Nhel*I at the beginning and *Xho*I at the end, this sequence could be cloned in the PYD1 plasmid as an expression vector in *Saccharomyces cerevisiae (EBY100)*. The virtual cloning of this chimeric sequence was performed and the results is shown in Figure 5.

**Figure 4.**
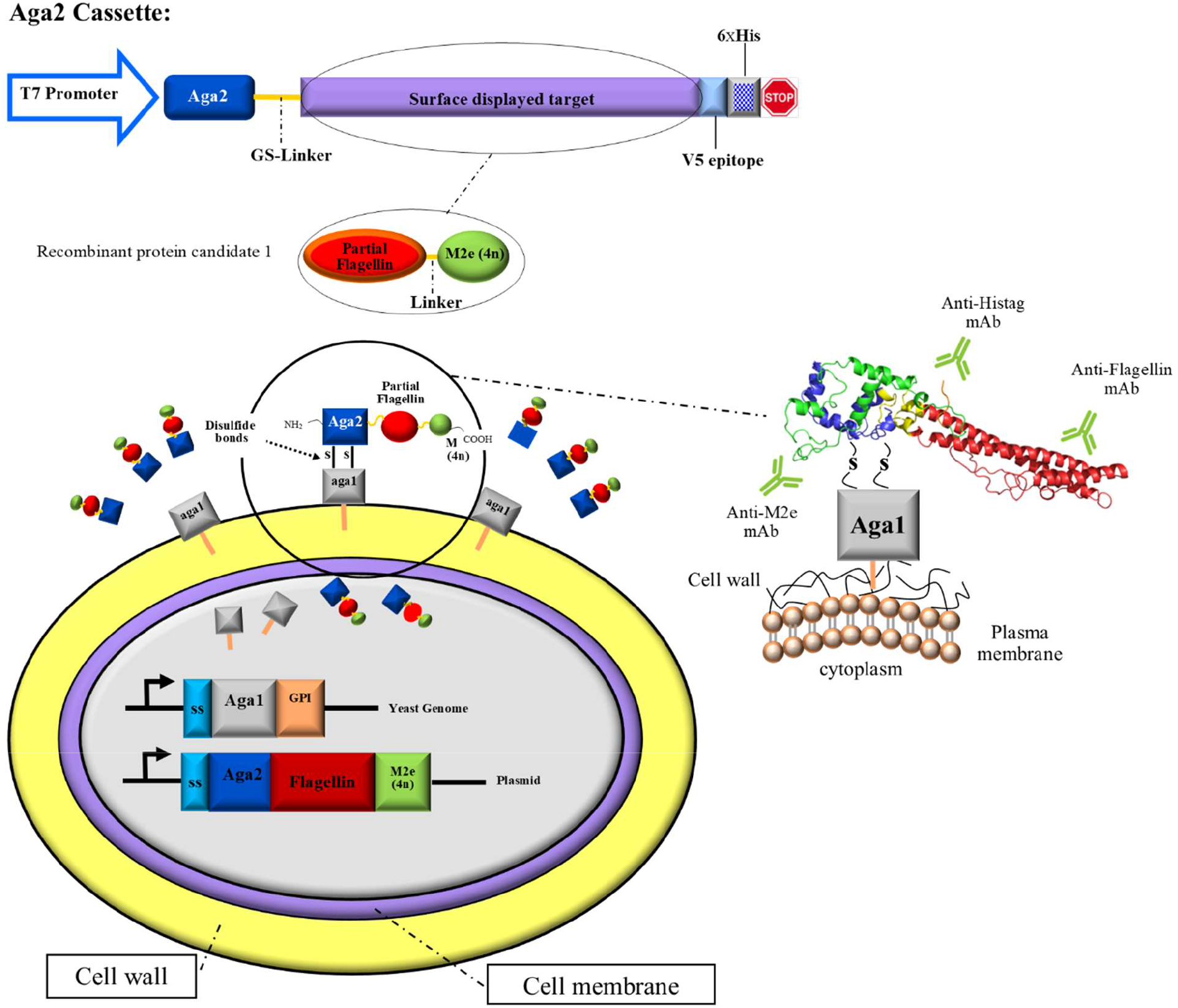
**a**) The Aga2 cassette consists of the target sequence which will be surface-displayed flanked by V5 epitope on the right side for detection purposes and Aga2 and linker on the left side for surface-displaying purposes. **b)** In the present study, the target sequence is M2e(4n) fused to partial sequence of flagellin by a linker. **c)** After expression of host genome (carrying Aga1 genome) and PYD1 vector (carrying the recombinant protein candidate 1), GPI anchor in yeast genome prevents the release of Aga1 into the media. Accordingly, the Aga1 would be connected to the cell wall through covalent bonds. Recombinant protein candidate 1 would be released to the media because of the signal peptide. Afterwards, the Aga1 and Aga2 make two disulfide bonds which will cause the surface-displaying of recombinant candidate 1. The result is different detectable targets for humoral immunity including His-tag for detection purposes, Flagellin as an adjuvant and M2e (4n) as the main antigen to create the immunity against variety of influenza subtypes.

**Figure 5.**
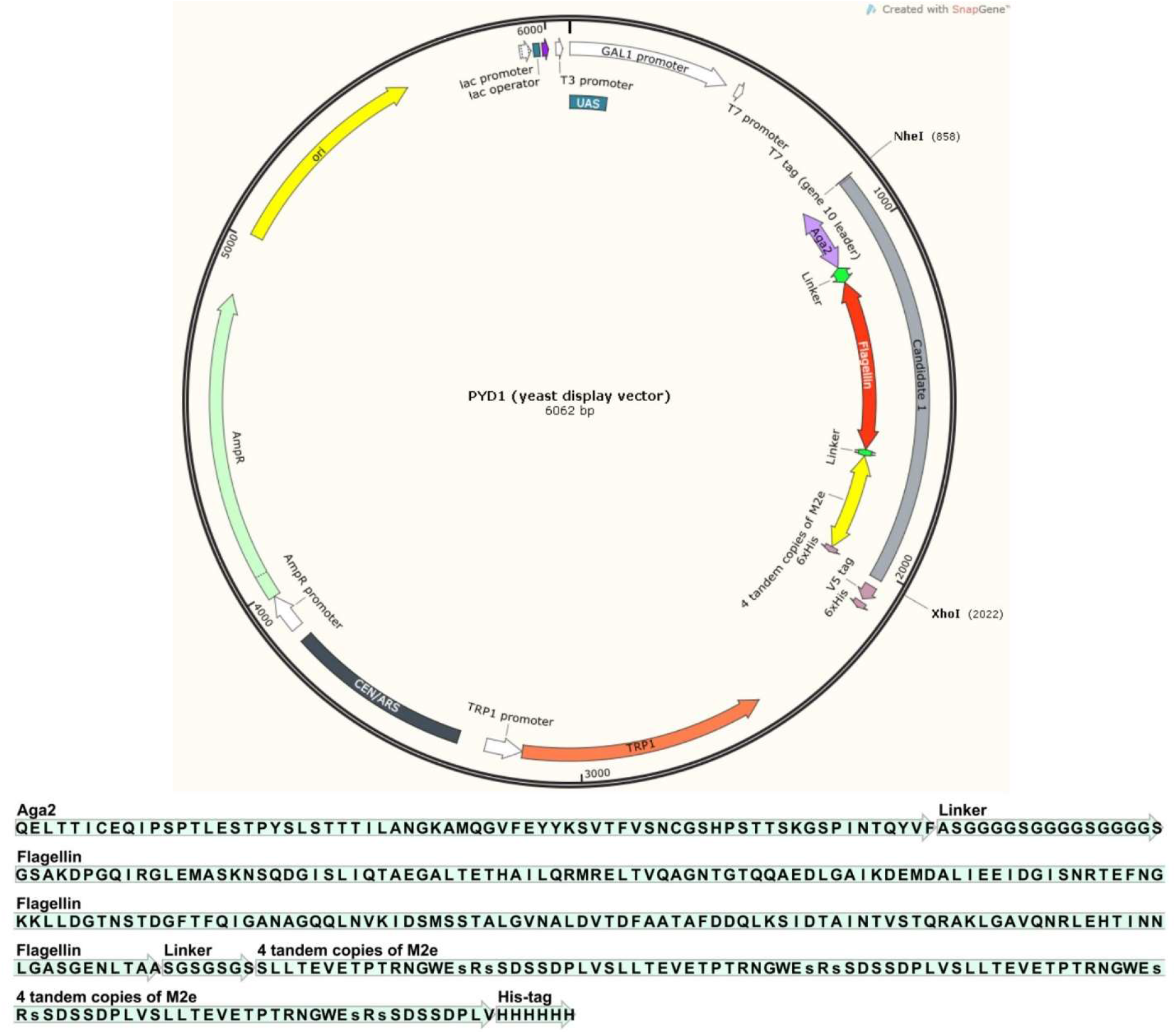
Virtual cloning of candidate 1 chimeric protein in PYD1 expression vector.

## Discussion

The pharmaceutical industry has been a significant respected industry player since 19^th^ century when it was first organized and churning out life-saving drugs such as penicillin. Since then, many developments have occurred such as vaccine design and production. Therefore, in essence this sector has been associated with saving lives and thus has been held in high esteem. But now, with the high cost of drug discovery and vaccine development, time-consuming process and probable adverse side effects, and on top of them, more awareness and education on the part of the consumers, the industry image has been battered. So, it is imperative that the sector finds ways of reducing the expenditures and time of drug discovery and vaccine development (such as drug repositioning (70, 71)) and escalating efficiency. Bioinformatics is one of the tools the industry has recently engaged to aid in the drug discovery (72-74) and vaccine development process (75-79) as well as to cut costs and the timelines. Accordingly, in this study we aimed to design and analyze the probability of producing a universal and cross-protective avian influenza vaccine through bioinformatic tools to diminish unnecessary laboratory costs.

In order to produce potential vaccines for various subtypes of influenza, many efforts have been done so far. These efforts have failed due to high mutation rate of HA and NA as two major antigens of influenza virus(80). Hence, it is rational to consider a preserved antigen against mutations to produce a cross-protective vaccine against various subtypes of influenza. M2e protein is one of the substantial antigens in this field. M2e peptide sequence has remained remarkably unchanged in influenza type A isolated since 1918 (13, 81) and this feature made it an appropriate target for vaccine design. Based on previous studies, evoked antibodies against the epitopes of M2e antigen have diminished the growth of influenza virus in *in vivo* and *in vitro* studies and created cross-reactive resistance to influenza A subtypes. Based on a study conducted by Huleatt et al. in 2008, among 7 sequences of M2e which was retrieved from 4 subtypes of influenza virus (H1N1, H2N2, H3N1, H3N2), a consensus sequence was obtained to be expressed in *E. coli*. Subsequently, mice immunized with this recombinant protein in aqueous buffer, without adjuvants or other formulation additives, developed potent M2e-specific antibody response. According to the importance of avian influenza disease in China, Japan, South Korea, Vietnam and India, 31 M2e protein sequences for H9N2 (12 sequences), H5N8 (6 sequences), H5N1 (7 sequences) and H7N9 (6 sequences) subtypes were extracted from NCBI database. The selected subtypes of influenza are the most important and the most damaging influenza subtypes in these countries. H9N2 inflicts widespread damage in Iran and Iraq every year (82).

In previous studies, a recombinant protein comprising the TLR5 ligand flagellin fused to four tandem copies of the ectodomain of the conserved influenza matrix protein M2 (M2e, to overcome the low antigenicity of M2e in comparison with HA and NA) was expressed in *E. Coli* and purified for homogeneity. This protein, retains TLR5 activity and exposes the epitopes of M2e which are identifiable by a specific monoclonal antibody, 14C2 (RCSB protein ID) (16). According to the purification costs and toxicity of components of *E. coli*, choosing a safe expression host seems to be advantageous. Recently, yeast-based vaccines have been investigated for influenza vaccination. Yeast as an expression host can facilitate and escalate the production of newly engineered antigens (45, 46).

First time, Li and colleagues in 2016 indicated that the yeast which express H5N1 Hemagglutinin at its surface can be used as an influenza vaccine. The reason of choosing yeast is its ability to perform post translational modifications and capability of stimulating immune system. This feature effectively activates dendritic cells and cytotoxic T cells (42). The recombinant yeast cells, simultaneously stimulate humoral and cellular immunity by presenting antigens to MHCI and MHCII pathways (44).

The same as previous study conducted by Huleatt et al. in 2008, (16) in our *in-silico* investigation 4 tandem copies of the obtained consensus sequences of M2e protein and a partial sequence of flagellin as an adjuvant were used to virtually be expressed at the surface of yeast. In this regard, two different structures were designed and their performance and features were investigated. Their features have been evaluated in various point of views. According to docking studies, the first candidate effectively expose its epitopes to a specific monoclonal antibody (MAB148). The complement-determining regions (CDRs) of MAB148 form a deep and narrow binding pocket that accommodates the N-Terminal part of M2e. Pro10 and Ile11 of M2e appear from the MAb148 binding pocket, where Pro10 kinks the M2e peptide in a way that its C-terminal segment is projected away from the monoclonal antibody (50). The two out of four epitopes of the first candidate resemble a fishing hook, with residues Ser2-Leu3-Leu4-Thr5-Glu6 forming a β-turn that is complementary to the shape of MAb148 paratope.

According to salleh et al. in 2012, by raising the compactness of protein, the stability will be increased subsequently. On the other hand, in spite of more sustainability of the second vaccine candidate, the epitopes of M2e antigen in this protein are covered by other sections and become undetectable by CDR region of monoclonal antibody. Accordingly, compactness of protein may have benefits and drawbacks simultaneously. It is predictable that despite greater stability of candidate 2 in comparison with candidate 1, this protein may evoke weaker immune response. According to MD simulation studies, Van der Waal, electrostatic and binding energies of “SLLTEVETP” epitope to the CDR region of MAb148 monoclonal antibody in the first design (the most important epitope of M2e (11, 50)) were considerably higher than second design. In addition, these estimations in the first design were significantly closer to positive control (crystallographic structure between “SLLTEVETP” epitope and MAB148 antibody) in comparison with candidate 2. As a result, contrary to the second candidate, first design may be able to provoke stronger humoral immune response. The importance of recognizing “SLLTEVETP” epitope has been discussed in previous studies. Cho et al. in 2016 (83) proved that serine 2, leucine 3, leucine 4 and threonine 5 are essential for binding to monoclonal antibody MAB148 of M2e protein. On the other hand, Grandea et al. in 2010 (12) represented that serine 2, threonine 5 and glutamic acid 6 are vital for binding to TCN-031 and TCN-032 monoclonal antibodies. They expressed that these antibodies can recognize a core in “SLLTE” section of N-Terminal region of M2e protein, which comprises amino acids 2 to 6. Therefore first candidate is expected to provide humoral immunity by MAB148, TCN-031 and TCN-032 antibodies. According to two exposed repetitions of “SLLTEVETP” epitope in the first design, there is a significant possibility of evoking three different specific monoclonal antibodies against this region.

PYD1 shuttle vector and *EBY100* yeast strain can be used to yeast-display the first candidate. pYD1 is a 5.0 kb expression vector designed for expression, secretion, and display of proteins on the extracellular surface of recombinant *Saccharomyces cerevisiae* cells (*EBY100*). Features of this vector allow regulated expression, secretion, and detection of expressed proteins on the cell surface of *EBY100* (84). The vector contains AGA2 gene from *Saccharomyces cerevisiae*. This gene encodes one of the subunits of the a-agglutinin receptor. Fusion of the gene of interest to AGA2 allows secretion and display of the protein of interest. *EBY100* expresses the *AGA1* gene under control of the *GAL1* promoter (84) and the attachment of AGA2 and AGA1 lead to yeast-displaying a recombinant protein.

In conclusion, the results of the present study propose a novel chimeric antigen which can be considered as a universal and cross-protective avian influenza vaccine candidate or as a complement of conventional avian influenza vaccines as well. In this regards, our laboratory has already initiated research in this direction.

## Acknowledgements

We would like to acknowledge Ferdowsi University of Mashhad for supporting this project.

## Declaration

### Authors approval

all authors have seen and approved the manuscript, and that it hasn’t been accepted or published elsewhere.

## Abbreviations

MD: Molecular dynamic
CDR: Complementarity-determining region
HA: Hemagglutinin
NA: Neuraminidase
M2e: Matrix 2 ectodomain
NCBI: National Center for Biotechnology Information
RCSB: The Research Collaboratory for Structural Bioinformatics
MHC: Major Histocompatibility Complex

